# Cell-type-specific circadian and light-responsive transcriptional dynamics in adult Drosophila neurons

**DOI:** 10.64898/2026.04.07.717038

**Authors:** Gillian Berglund, Pranav Ojha, Maria Ivanova, Melina Pérez Torres, Michael Rosbash

## Abstract

The *Drosophila* adult central brain contains 240 circadian neurons, of which there are more than 25 different neuron subtypes based on connectomic data. Recent single cell RNA-seq (scRNAseq) characterization of these neurons “around the clock” also indicates a similar number of molecular subtypes of circadian neurons, but other conclusions from these transcriptomic studies warranted verifying and extending with other approaches. To this end: 1) We used a genetic multiplexing strategy to profile the transcriptomes of circadian neurons from multiple time points in a single experiment, reducing confounding technical variation between timepoints; 2) Large numbers of single nuclei were sequenced (snRNA-seq), which was enabled because the new method EL-INTACT purifies nuclei from frozen heads; 3) We assayed 12 time points under both light–dark (LD) and constant darkness (DD) conditions. These approaches showed dramatic transcriptional differences between time points in many circadian neuron types and enhanced time-of-day gene expression analysis. The data indicate that most of this regulation is transcriptional and circadian. There were however a small number of light-dependent transcripts, including a few that correspond to mammalian immediate-early genes. They probably play a role in the light-regulation of gene expression and behavior in specific neurons, perhaps circadian entrainment or phase-shifting. The results taken together provide a more comprehensive picture of gene expression heterogeneity within adult *Drosophila* circadian neurons including how intrinsic clock mechanisms and light cues are integrated across circadian neuron subtypes.

## Introduction

*Drosophila melanogaster* is a powerful system for studying the cellular and molecular organization of the brain. Its sophisticated genetic toolkit, combined with a compact and numerically tractable nervous system, enables precise access to and characterization of neural circuits that underly a rich repertoire of complex and highly coordinated behaviors. Recent advances in high-resolution whole-brain connectomics (1, 2) and single-cell transcriptomic profiling (3–5) have further transformed the field. They reveal striking heterogeneity across the fly brain, including within anatomically defined neuronal populations (6–8). These findings indicate that many behaviorally relevant circuits in the *Drosophila* central brain are composed of many molecularly distinct neuron types with few copies/brain. This description also applies extremely well to the 240 circadian neurons of the central brain. There are at least 27 transcriptomic circadian cell types. Interestingly, this indicates that the fly circadian system may be much more heterogeneous that the 20,000 neurons of the mammalian brain pacemaker, the SCN (9).

This description of the fly circadian system is based on our scRNA-seq clock neuron transcriptome characterization, which was also done “around the clock” to profile the circadian regulation of gene expression. Because this regulation is known to be substantially transcriptional, we predicted snRNA-seq profiling might provide a view that is less influenced by post-transcriptional events. In addition, our previous scRNA-seq transcriptomic characterization of circadian neurons had an issue that we thought might be ameliorated by snRNA-seq profiling. Prominent circadian neurons, the famous large LNs, were almost entirely absent, and other important large neurosecretory circadian neurons were poorly recovered in the 10X protocol (6, 10); this should be much less problematic with nuclear purification.

We were also encouraged in this direction by a recent nuclear purification method developed in our lab, EL-INTACT (11). Although it was designed for chromatin studies of small numbers of discrete fly brain neurons, it was easily optimized for snRNA-Seq. (See Methods.) EL-INTACT completely solved the missing clock neuron subtype problem in our previous 10X cell data. More importantly, EL-INTACT begins with large numbers of frozen fly heads rather than dissected brains, which allowed a remarkable coverage of rare neurons that are otherwise very difficult to collect for transcriptomic studies. A second improvement over our previous data (and all existing circadian gene expression data) was the use of a genetic multiplexing strategy (12), in which chromosomes from wild-type strains of the *Drosophila* Genetic Reference Panel (DGRP) as molecular tags, served as molecular tags (13), allowing profiling of multiple circadian time points together and reducing confounding technical variation in circadian gene expression analysis (13, 14).

These new strategies together with assays in constant darkness (DD) as well as light-dark (LD) conditions combined with the large number of cycling transcripts within many circadian neurons to reveal remarkable transcriptomic heterogeneity even within single cell types. Importantly, these new results establish a more comprehensive, cell-type-resolved view of circadian and light-responsive transcription in adult *Drosophila* clock neurons, which now provide in turn a better platform for investigating the integration of neuron type-, circadian-, and light-regulation of clock neuron gene expression and behavior.

## Results

### Exploiting El-INTACT and DGRP chromosomes to profile single-nuclei RNA-Seq of *Drosophila* circadian neurons

As described above, we sought to overcome two technical obstacles that interfere with optimizing single nuclei transcriptomic data from *Drosophila* adult circadian neurons. The first is collecting sufficient high quality nuclei from these quite rare neurons for snRNA-Seq. To this end, we modified the nuclear purification method EL-INTACT to improve the yield and reduce the time necessary for isolation. EL-INTACT was originally designed for chromatin studies and purifies nuclei via a FLAG tag on the nuclear surface followed by fluorescence-activated cell sorting (FACS) (Fig. S1A) (11).

The second is confounding technical variability (batch effects) that is inherent to comparing scRNA-seq datasets that were generated in separate experiments; it is particularly problematic for circadian time point comparisons. Which differences are biological, namely gene expression differences at different times of day, and which differences are due to batch effects?

To address this issue genetic multiplexing strategy, previously used for pooled profiling of Drosophila neurons across multiple stages of development (14). Samples from different circadian time points and replicates were tagged using wild-type chromosomes from the *Drosophila* Genetic Reference Panel (DGRP) strains (13, 15). This approach allows the profiling of multiple time points within a single experiment and sample, thereby minimizing batch effects and strengthening comparisons. Flies were entrained to a 12:12 light: dark (LD) cycle for at least 3 days, and 6 timepoints were processed simultaneously. We performed this twice, each covering 6 timepoints separated by 4 hours, with two independent DGRP genotypes for each time point as biological replicates. The two datasets were shifted by two hours to generate a combined circadian dataset of 12 time points, spaced evenly every 2 hours around the clock (Fig S1A). For each dataset, we sequenced 9,312 and 9,835 nuclei from each; because of EL-INTACT, this is considerably more than what had been sequenced in our prior circadian scRNA-seq experiments (6, 7, 10).

Applying unbiased filtering and removing doublets (see Methods for the filtering cutoffs) resulted in a high confidence single nuclei dataset. It was then combined with our previous scRNA-seq datasets of circadian neurons, and we then used an unsupervised clustering algorithm to integrate the two datasets and assign cell identities to the clusters. This resulted in 24 circadian clusters containing 9,899 circadian nuclei from the 12 timepoints. The clusters included all previously reported circadian cell types with the exception of some missing DN3 clusters (Figure 1A, left), almost certainly for simple differences between the studies in driver-expression patterns. Importantly, the nuclei contain proper numbers of three cell types that were underrepresented in past 10X cell datasets (Fig. S1B). (See Discussion.)

**Fig. 1.**
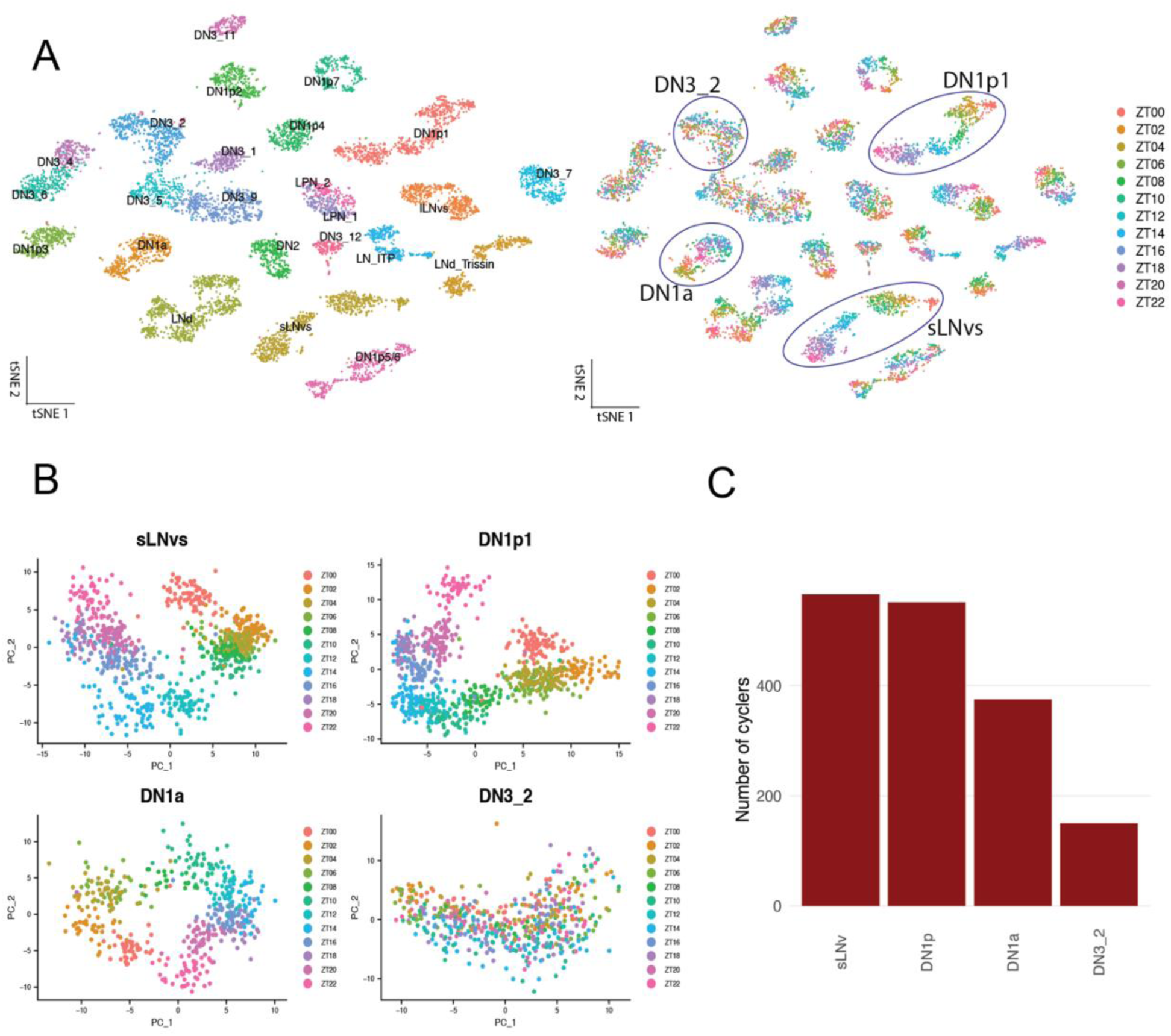
snRNA-Seq of *Drosophila* circadian neurons reveal timepoint separation and neuron-selective phenotypes. A) T-distributed stochastic neighbor embedding (t-SNE) plot showing the 9,899 circadian nuclei grouped into 24 clusters, colored by cell type (left) and circadian timepoint (right). B) Principal component analysis (PCA) of subclustered sLNvs, DN1p1s, DN1a and DN3_2 cell types with 814, 807, 471 and 550 nuclei, respectively. C) Number of cycling transcripts identified in each cell type plotted in B. Cycling transcripts defined by a JTK_cycle P value of less than 0.05.

Many of these circadian clusters have strikingly spread-out patterns in the t-Distributed Stochastic Neighbor Embedding (t-SNE) plots (Fig. 1A). They likely reflect distinct nuclear transcriptomes at different time points as well as the deep dataset, an average of 300-400 nuclei/cluster or >30 nuclei/time point/cluster. The coloring of the individual time points strongly supports this interpretation: Nuclei of one color (from one time point) are all together, and adjacent time points/colors are adjacent within the cluster, e.g., ZT8 is next to ZT10; in contrast, ZT8 and ZT20 are far apart within the cluster as expected for time points that are 12hrs different. Importantly, this heterogeneity is likely driven by biological differences between time points as they are all pooled and profiled in the same experiment.

Some cell types like the small LNvs and DN1p1s displayed this spread-out pattern clearly, whereas others like a subtype of DN3 neurons not at all (Fig. 1A). As the interpretation of tSNE plots of scRNA-seq datasets can be challenging, we performed separate Principal Component Analysis (PCA) on individual clusters, including sLNv, DN1p1, DN1a, and DN3_2 (Fig. 1B). The first two principal components (PC1 and PC2) separated three of these clusters by time points (with the exception of DN3_2). Cells from adjacent time points were adjacent in PCA space, suggesting that this heterogeneity reflects temporal changes in transcriptomes across circadian time points. Based on previous single cell data (10), cell types with less distinct timepoint patterning like the DN3 subtype could be due to less gene expression differences between timepoints. To this end, we used JTK_CYCLE to compare the number of cycling genes identified in the four sampled cell types. The number of cycling transcripts with a JTK pvalue of less than 0.05 was consistent with previous findings (10): The sLNvs and DN1p1s had 566 and 551 genes identified respectively, whereas the DN3 subtype had nearly 75% fewer cycling genes with only 150 gene identified (Figure 1C).

### How do the patterns change in DD (constant darkness)?

To assay the effect of light, i.e., to test whether the striking time point separation of many circadian clusters is under circadian and/or light-regulation, we sequenced clock neuron nuclei on day 2 of constant darkness (DD). We followed the same experimental design as under LD conditions, namely two experimental replicates and 12 time points. A total of 21,197 nuclei were sequenced, and the final DD dataset contained 11,256 nuclei following filtering and identification of circadian nuclei.

There was very similar time-of-day separation within clusters and the same variability between circadian cell types in DD as in LD (Fig. 2A). Moreover, core clock gene expression is very similar under the two conditions; *tim* is an exception with obvious lower amplitude cycling in DD than in LD (Fig. 2B, Fig S2). (See Discussion for more detail.) Another exception is the large LNv cluster, which has little or no core clock gene expression and mRNA cycling in DD; this is consistent with previous findings (16).

**Fig. 2.**
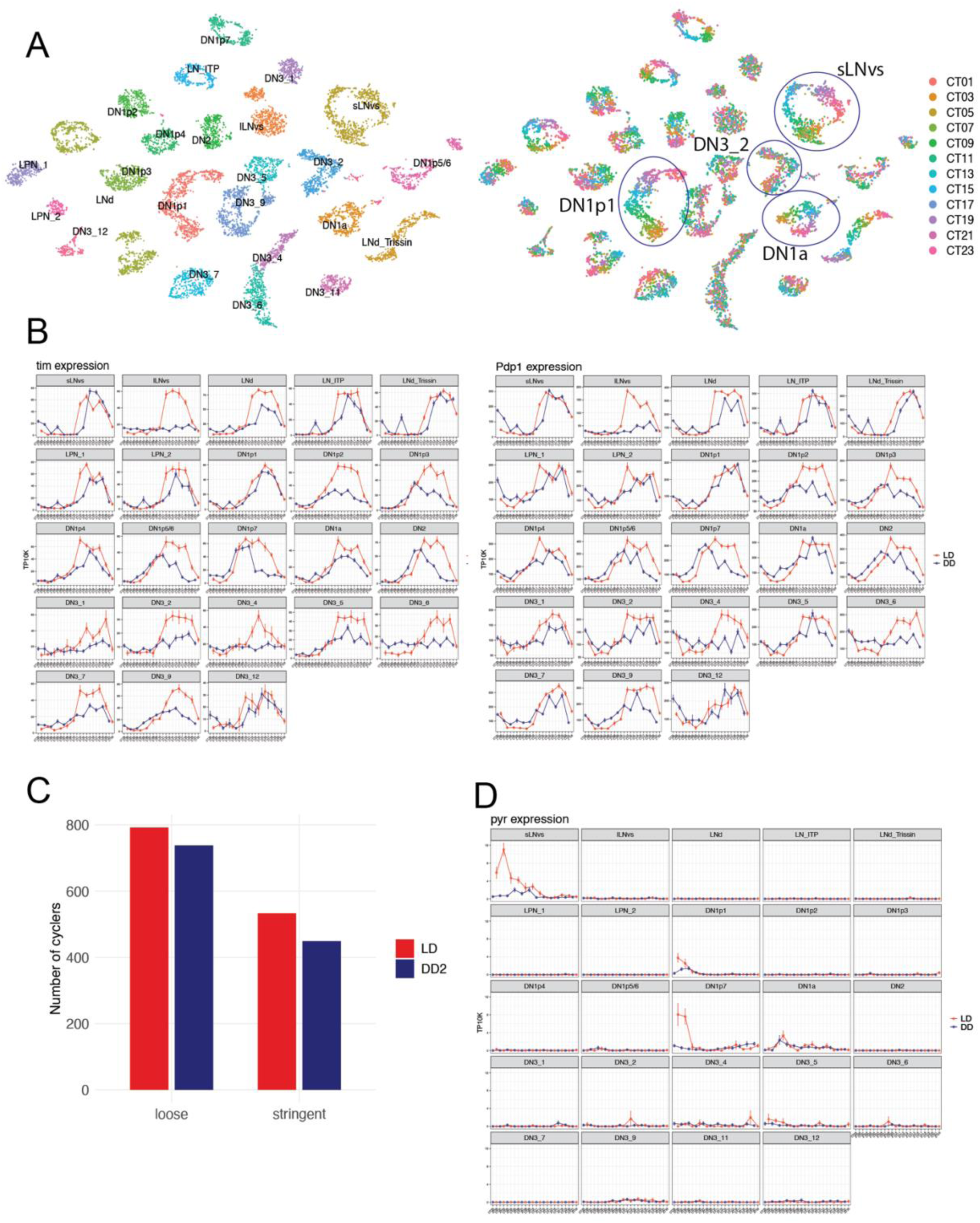
snRNA-seq of *Drosophila* circadian neurons in the second day of DD maintains timepoint separation and cycling gene expression. A) t-SNE of 9,835 circadian nuclei collected across 12 timepoints in the second day of constant darkness. Clusters are colored by cell type (left) and circadian timepoint (right). B) The mean *tim*, and *Pdp1* expression throughout the day in light: dark (LD) condition (red) and constant dark (DD) (blue) is graphed for each cluster. Error bars represent mean ± SEM. C) Number cycling genes identified in LD and DD under different cycling cutoffs. Loose cutoff: JTK p-value less than 0.05 Stringent cutoff: JTK p-value less than 0.05, a maximal expression of at least 0.5 TP10k, and a cycling amplitude (maximum expression divided by minimum expression) of at least twofold. D) The mean *pyr* expression throughout the day in light: dark (LD) condition (red) and constant dark (DD) in blue is graphed for each cluster. Error bars represent mean ± SEM

Moreover, the number of LD cycling transcripts is similar in DD, defined under more stringent as well as more relaxed cycling criteria (Fig. 2C). The few notable exceptions are almost all cell type-specific (Fig. 2D and Discussion). Not surprisingly, the large LNvs contain many cycling transcripts in LD and essentially none in DD, consistent with its core clock gene expression.

### How do the light transitions (lights-on and lights-off) influence gene expression?

The above LD vs DD description and data ignore a key difference between the two conditions: a few genes are potently expressed only at the single ZT0 time point (Fig. 3A). Relevant here is that the ZT0 time point is taken 15-20 min after the lights turn on, i.e., not precisely at ZT0, suggesting that these genes are light-activated. Two of these ZT0 genes are *Hr38* and *sr*. These genes and others were previously identified as activity-regulated genes (ARGs) in *Drosophila* circadian neurons and elsewhere (17). This lights-on response is neuron-specific as LNvs respond strongly whereas many other circadian neurons show little or no *Hr38* and *sr* activity at ZT0. The LNvs response is a sharp burst of *Hr38* and *sr* expression; there is essentially no detectable expression at ZT22 or at ZT2, 2 hours before and 2 hours after the lights-on event, as well no expression at CT0 (Fig. 3A).

**Fig. 3.**
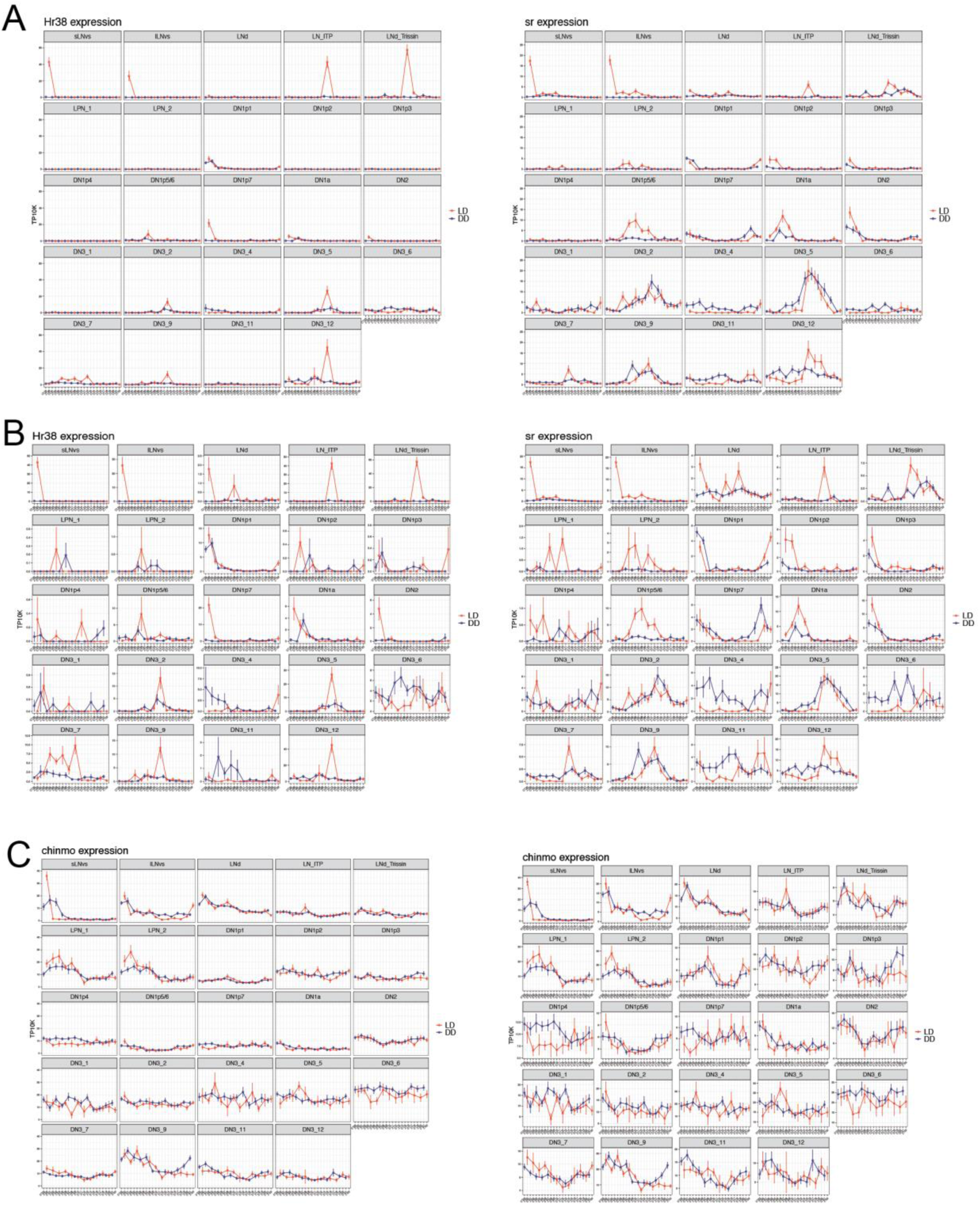
Neuron specific expression of *Drosophila* activity-regulated genes in response to light stimulation A) *Hr38* and *Sr* expression throughout the day in light:dark (LD) condition and constant dark (DD) condition for each cluster. X axis indicates the time points in LD and DD. B) *Hr38* and *Sr* expression throughout the day in light:dark (LD) condition and constant dark (DD) condition with variable Y-axes scales for each cell type. C) *Chinmo* expression throughout the day in light:dark (LD) condition and constant dark (DD) condition with a fixed y axis (left) and variable Y-axes ranges (right) for each cluster. Red and blue lines indicate the gene expression in LD and DD conditions, respectively. Error bars represent mean ± SEM.

To address this time course in more detail, we isolated circadian nuclei 15 min, 30 min and 1hour after lights-on and the same circadian timepoint without turning lights on using the experimental method and design as described above (two DGPR genotypes per condition). *Hr38* and *sr* transcript levels peak at 15-30 minutes following lights-on and have substantially decreased by ZT1, one hour later (Fig. S3). This time course mimics a canonical IEG expression pattern and suggests that lights-on stimulates neuronal activity in the LNvs, which then transiently upregulates IEG/ARG gene expression.

Notably, other circadian cell types also show increased *Hr38* expression at ZT0, including some DN1ps and the DN1as. Although a simple Y axis scale change makes light-mediated expression in these neurons unambiguous, it is much less dramatic than in the LNvs, which have a log fold change of 8.03 and 7.25 for *Hr38* in the small and large LNvs, respectively. Another difference is that the DN1p1 and DN1a cells but not the LNvs manifest considerable *Hr38* expression even in constant darkness (Fig. 3A). *Sr* is also expressed beyond the LNvs, but these patterns are even more complicated than those of *Hr38*, making it difficult to cleanly distinguish between light and circadian regulation within these neuron types (See Discussion).

*Hr38* as well as *sr* manifest a second burst of expression. It is at ZT12 (lights-off) and in different neurons from the LNvs: this ZT12 expression is in the ITP- and Trissin-expressing LNds, which are two of the three classes of evening cells. (Note: ZT12 was really ZT12.33, i.e., 20 min after lights-off.) This *sr* ZT12 pattern is less striking as well as more complicated than the *Hr38* ZT12 pattern (*sr* has considerable DD expression), but lights-off clearly triggers or enhances the expression of both genes (Fig. 3A). This lights-off evening cell stimulation is also more complicated to interpret than the lights-on LNv stimulation but likely reflects disinhibition: neuronal activity in these two evening cells is presumably inhibited by the constant presence of light between ZT0 and ZT12 and is relieved by lights-off at ZT12 resulting in ARG expression. (See Discussion.)

*Hr38* and *sr* are the most well-known *Drosophila* ARGs/IEGs, but our original circadian neuron stimulation work (17) identified additional candidate ARGs including *Chinmo* (Fig. 3C). Its patterns are interesting: The *Chinmo* s-LNv pattern appears light-stimulated similar to that of *Hr38*, but the *Chinmo* patterns in other neurons are very similar between DD and LD conditions. Surprisingly, these patterns even include the l-LNvs, which are not supposed to have cycling transcripts. The similar DD and LD patterns may indicate that *Chinmo*-rather than *Hr38*- or *sr*-transcription mirrors circadian neuronal activity.

### How do these nuclear RNA profiles differ from cell (neuron) RNA profiles?

Which features of these nuclear data are also present or absent in data from whole cells, in this case intact neurons? The answers to this question may have implications for circadian regulation, for example transcriptional vs post-transcriptional regulation. One striking feature of the snRNA-seq dataset is its timepoint separation (Fig. 1A), which has not been described in previous circadian neuron datasets. To address whether this feature is restricted to the snRNA-seq DGRP data, we compared our data with scRNA-seq DGRP data generated with the same CLK856-GAL4 driver (18) (Fig. 4A). Although there are only six time points in the cell data rather than the twelve in Fig. 1, it is evident from the colored cells that the adjacent time points subcluster together similarly within many clusters (Fig. 4A) like in the nuclear RNA-Seq pattern (Fig. 1A). Although this experiment does not show that the two patterns are indistinguishable, i.e., that there are no differences between snRNAseq and scRNAseq, it does indicate that biological time point separation is not unique to nuclei.

**Fig. 4.**
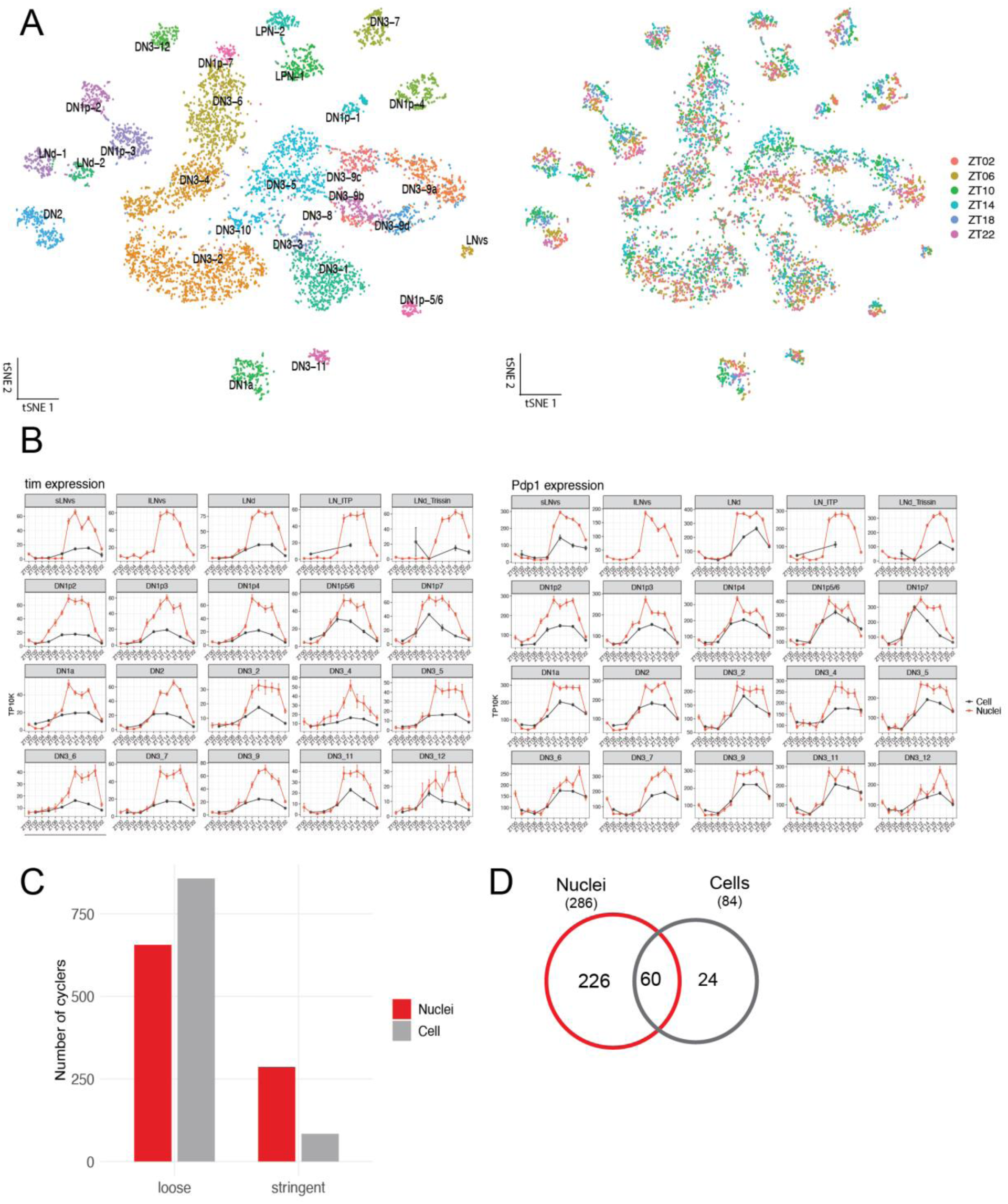
Comparison of scRNAseq and snRNAseq circadian datasets A) t-SNE of circadian scRNAseq collected across 6 timepoints of LD. Clusters are colored by cell type (left) and circadian timepoint (right). B) *Timeless* and *Pdp1* expression throughout the day in light:dark (LD) condition for each cluster. Red and black lines indicate the gene expression in nuclei and cell datasets, respectively. Error bars represent mean ± SEM. C) Number cycling genes identified in cell and nuclei datasets with different cycling cutoffs. Loose cutoff: JTK p-value less than 0.05 Stringent cutoff: JTK p-value less than 0.05, a maximal expression of at least 0.5 TP10k, and a cycling amplitude (maximum expression divided by minimum expression) of at least twofold. D) Comparison of cycling genes identified in nuclei and cell datasets.

A second comparison between nuclei and cells is the cycling patterns of the core clock transcripts within individual circadian cell types (Fig. 4B, S4). These patterns have some differences between cell types/clusters but are otherwise quite similar. There is however one feature that is apparent for all clock genes and most if not all circadian clusters: the cycling amplitude appears considerably higher for nuclear RNA than for cell RNA; this is especially prominent for *tim* RNA. Although this difference could be influenced by differences between the two normalization processes, cells vs nuclei, and the plots reflect 6 time points for the cell data vs the 12 time points comparisons available for the intra-nuclei data, there were similar, major differences in RNA amplitude fold-change: nuclear peak:trough ratios >> cytoplasmic peak:trough ratios, indicating that the cycling amplitude of the core clock genes is indeed higher in nuclei than in cells (Fig. 4B, S4).

To address the extent to which these nuclei vs cell differences are general and not specific to core circadian genes, we extended this comparison to all cycling genes within the circadian neurons, i.e., are there many more cycling genes in one compartment than the other (Fig. 4C)? Because we never generated 12 time points of cell data (and because the number of time points has a major influence on the number of identified cycling genes), we used the same 6 time points of nuclear DGRP data to compare with the 6 time points of cell DGRP data. Although 6 time points is a sub-optimal cycling criterion (19), it should be sufficient to reveal substantial differences between nuclear and cell data.

Under fairly loose cycling criteria, the number of cycling transcripts is quite comparable between the cell and the nuclear datasets, no more than 25% greater in the cell data than in the nuclear data. Under more stringent criteria however, the difference is much more dramatic and reversed: There are more than 3X as many cyclers from the nuclear dataset than from the cytoplasmic dataset (Fig. 4C). The most important stringent criterion addition is a two-fold amplitude requirement. This suggests that there are many more higher amplitude nuclear cyclers than cytoplasmic cyclers, which is verified by inspecting the plots of individual genes (Fig. S4); a cycling amplitude difference is similar to the conclusion from the core clock transcript cycling curve comparisons (Fig. 4B). Perhaps RNA turnover in the cytoplasm dampens the amplitude of circadian transcriptional regulation. (See Discussion.)

## Discussion

Our previous scRNA-seq transcriptomic characterization of *Drosophila* circadian neurons was less than optimal for a number of reasons. 1) Temporal analysis was compromised by possible technical variation due to the separate processing of each time point 2) Specific known neurons were poorly recovered. 3) Light-regulated transcripts were not detected. 4) Possible post-transcriptional vs transcriptional regulation of mRNA cycling was not addressed. To improve this characterization, we sequenced nuclei from almost the same biological material but with two important technical modifications: We optimized the chromatin purification method EL-INTACT (11) for snRNA-seq, and used the genetic multiplexing strategy (12, 14) to process circadian time points together and eliminate batch effects. We also assayed a denser series of 12 evenly spaced time points under both light–dark (LD) and constant darkness (DD) conditions. EL-INTACT recovered many more neurons than traditional protocols, especially previously problematic neurons, and together with the pooled profiling of all time points created a superior circadian dataset.

Our previous single cell circadian neuron sequencing efforts began with a cell-seq assay and then transitioned to droplet-based 10X Genomics assays (6, 7, 10). Using previously assigned marker genes, all droplet-based experiments were almost entirely missing the l-LNvs; yet their s-LNv neighbors were present as expected. As the CLK856 driver expresses similarly in both neuron types and they both are present at 8 cells/brain, the famous l-LNvs should have been recovered. In addition, the trissin and ITP-expressing LNds were recovered but were underrepresented relative to their in vivo cell numbers. As all three of these are large neurosecretory neurons, they likely present a similar recovery issue during some step of the 10X Genomics protocol. This problem is absent from the snRNA-seq data, probably because their nuclei are not very different from nuclei of smaller neurons (20). The only other cell type identification difference with our previous neuron data is that many DN3 neuron clusters are absent in these new nuclear data. This is almost certainly because of reporter gene expression differences: our most recent clock neuron sequencing work used a dedicated DN3 driver in addition to CLK856 as well as a different reporter gene from the one used here (10), which almost certainly favored DN3 expression relative to this nuclear sequencing effort.

These exceptions notwithstanding, cell type identification in this work matched very well our previous cell type identification based on scRNA-seq (Fig. 1A). Notably, many of the individual clusters in this study display striking time-of-day heterogeneity; this substructure is likely due to their large number of cycling transcripts (Figs. 1D, 4C). Likely important as well are the substantial technical improvements: 1) the large number of nuclei sequenced, because of El-INTACT and its use of frozen heads rather than brain dissection; 2) the contribution of the pooled processing using the DGRP chromosomes; 3) the use of 12 rather than 6 time points. With respect to these last two factors, we note that the genetic multiplexing strategy can process 12 samples together. For logistical reasons however, we collected and processed them in two separate groups of 6: times 2, 6, 10, 14, 18, 22 and then times 0, 4, 8, 12, 16, 20. Although this arrangement could cause batch effects, important ones are unlikely because the two sets of time point data interdigitate smoothly (e.g., Fig. 2B).

The striking time-of-day heterogeneity was similarly apparent under DD conditions (Fig. 2A). These parallels fit with the generally modest effect of DD compared to LD, only a decrease in the amplitude of clock gene expression in DD relative to LD. This effect is most apparent for *tim*, which has for unknown reasons the biggest effect of light on circadian amplitude (Fig. 2B, Fig S2). A decreased amplitude of clock gene expression in DD is most easily seen in pseudobulk analysis, i.e., by assaying clock gene cycling in all clock neurons together; this decreased amplitude is also apparent by assaying fold-change, i.e., peak:trough values decrease markedly from LD to DD (data not shown).

Although there is not much difference in the number of cycling transcripts between LD and DD, there are a few exceptions, i.e., genes that appear to strictly require LD conditions for their clock neuron cycling. Most of them appear to be only expressed in a restricted number of clock neuron types (Fig. 2D). Also present in this category is the gene *Hr38*. It has the most striking qualitative difference between LD and DD, and its expression has been previously used to identify active neurons in insects (21–23) and quite recently in *Drosophila* (24). *Hr38* and the gene *sr* are the fly orthologs of genes previously identified as ARGs or IEGs in mammals, namely, the NR4A family of nuclear hormone receptors and Egr-1 (or Zif268), respectively. LNvs show a dramatic and rapid induction of *Hr38* shortly (20 min) after lights-on at ZT0. There is no expression in DD, indicating that it reflects a direct or indirect light-mediated response. As expression returns to baseline by ZT2, this likely explains why we never saw *Hr38 e*xpression in our previous studies; we always began our six time point fly harvestings with ZT 2 and ended with ZT22 (10). Notably, *Hr38* is also expressed at ZT0 in DN1a and DN1p1 neurons, but this transcript is present there in DD as well as LD; this indicates that its expression in these neurons is more complicated than just light-mediated. The expression of other ARGs/IEGs like *sr* and *chinmo* is also influenced by light, but these patterns are even more complicated than those of *Hr38.* More mechanistic insight into all of these LD vs DD differences await further experiments.

The differences between these snRNA-seq data and scRNA-seq data are less certain than those between LD and DD. This is because the experimental protocols for snRNA-seq and scRNA-seq are not identical. Moreover, there are only six time points from cells, which is suboptimal for cycling analysis (19). Nonetheless, rough comparisons suggest that clock gene cycling amplitude is greater in nuclei than in neurons (Fig. 4B). The data suggest that this is also true for most cycling transcripts, which also explains why there are more cycling genes in the nuclear data than in the cell data (Fig. 4C). A simple interpretation is that most circadian gene expression regulation is transcriptional and that cytoplasmic amplitude is decreased by post-transcriptional cytoplasmic mechanisms like RNA turnover. Exceptional cycling transcripts with greater cytoplasmic than nuclear amplitude might then have a more dominant post-transcriptional regulatory mechanism.

To what extent are these conclusions unique to clock neurons and to *Drosophila*? Might other cells, tissues and organisms with canonical molecular clocks give rise to the same conclusions? The striking intra-cell type heterogeneity probably depends on the extent to which these other cells and tissues harbor a large number of cycling genes beyond core clock genes; mammalian liver is a good candidate. Although higher cycling amplitude in nuclei than cells and the clock gene cycling amplitude difference between LD and DD have not been reported elsewhere, it may exist. Perhaps food, nutrition and chemical stimuli can mimic light in many other cells and tissues, similar to the role that ligands and receptors play in the stimulation of IEG expression.

In any case, the data here provide a comprehensive view of how cell type, intrinsic clock mechanism and environmental light cues are differentially integrated across circadian neuron subtypes in *Drosophila*. We initially thought that the large number of circadian cell types by molecular criteria might be exceptional. However, the extensive circadian neuron heterogeneity is also apparent morphologically (8). Moreover, the quite few cells in each neuron type/brain appears characteristic of most fly central brain neurons (5) and underscores the importance of neuron or nuclear purification for proper characterization. Otherwise, there is insufficient coverage, which explains why whole fly or brain scRNA-seq or snRNA-seq data show such little central brain detail (5). In contrast to the general nature of this transcriptomic heterogeneity, the striking intra-cluster heterogeneity of many circadian neuron cell types is probably less general as it likely relies on their large number of robust cycling transcripts (Fig. 1D). Perhaps other central brain neurons will show similar intra-cluster heterogeneity if their function can be exploited to assay the equivalent of time points, e.g., food content, satiety, level of aggression etc.

## Methods

### Fly stocks and rearing for single-cell RNA-seq experiments

*Drosophila* melanogaster stocks were housed in standard medium (cornmeal/agar/yeast) under 12h:12h light/dark cycle at 25°C. UAS-unc-84-tdtomato-FLAG flies were a gift from Sean Eddy. Female flies expressing the Clk856-Gal4 driver with the tdtomato-FLAG reporter (UAS-unc-84-tdtomato-FLAG) were crossed to male flies carrying one of twelve unique third chromosomes from the wild-type strains of the *Drosophila* Reference Genetic Panel (DGRP). These strains also carried UAS-His2A-GFP transgenes, which were not used in our experiments. This panel of DGRP-tagged strains was previously used for pooled profiling of the developing visual system neurons (14, 15). Flies with the same DGRP genotype were combined and entrained for at least 4 days in 12:12 LD. Two unique DGRP genotypes were assigned to each timepoint as biological replicates. Flies were collected at ZT02, ZT06, ZT10, ZT14, ZT18, and ZT22 for the first batch. DGRP lines were then reshuffled, and flies were entrained and collected at ZT00, ZT04, ZT08, ZT12, ZT16, and ZT20. For collection at ZT00 and ZT12, flies were collected 15 minutes after the lights switch on or off respectively, to minimize startle effects.

Approximately 1500-3000 flies were used for each single nuclei experiment. Single nuclei were isolated as previously described in Ojha et al. (11). In brief, fly heads were isolated with vigorous vortexing followed by separation over dry ice cooled sieves. 1500-3000 fly heads were added to 5 mL of homogenization buffer [250 mM sucrose, 1M Tris (pH 8.0), 1M KCl, 1M MgCL_2_, 10% Triton-X 100, 0.1M DTT 1X complete protease inhibitor cocktail]. After 20 tractions of the mini-homogenizer at 1000 rpm in two 15 ml dounce homogenizers, nuclei were filtered through 10 µm CellTrics strainer (Sysmex: 04-004-2324) and spun down at 800 g for 10 minutes at 4°C. Nuclei were resuspended in resuspension buffer (1% BSA in PBS with 0.1% Tween-20) and incubated with 40 μl of anti-FLAG magnetic beads (FisherSci, PIA36797) for 45 minutes at 4°C with constant end-over-end agitation. Bead-bound nuclei were then collected on a magnet and washed once with resuspension buffer. Flag beads were competitively eluted off with 3XFlag-peptide (Proteintech fp), shaking on a thermomixer for 10 minutes at 600 rpm. Flag beads were collected on a magnet and the supernatant containing isolated nuclei were filtered through a 35 μm cell strainer into a 5 mL round bottom polystyrene FACS tube. Hoechst dye (one drop per 0.5 ml of sample; Invitrogen, #R37605) was added into the sample tube to stain the nucleus.

To optimize El-INTACT for RNAseq, all steps were performed at 4°C and RNase inhibitors were added to homogenization and resuspension buffers at a 0.5% concentration. Importantly, the elution of nuclei with FLAG peptides requires using buffers without RNAse inhibitors due to reducing agents in RNAsin interfering with the peptide and reducing nuclei yield. All steps downstream of elution including FACS and preparing samples for 10X genomics use RNase inhibitors. A BD Melody FACS machine in single cell sorting mode was used for cell collection.

TdTomato- and Hoechst-positive single cells were collected in a 1.5-ml Eppendorf tube containing 0.4 ml of collection buffer [phosphate-buffered saline (PBS) + 1% bovine serum albumin + 0.5% RNasin] and used for downstream single-cell RNA sequencing library preparation.

### Library preparation and raw data processing

Isolated nuclei from FACS were spun down on a benchtop centrifuge at 1000g for 10 min at 4°C before loaded to the GEM chip from the Chromium GEM-X Single Cell 3′ Kit (version 4) of 10X Genomics. The libraries were prepared according to the standard user guide (CG000731 Rev. B) from 10X without any modifications. 10X libraries were sequenced by Illumina Nextseq 1000 with the P2 XLEAP-SBS kit (100 cycles) using a paired-end sequencing workflow and with recommended 10x v4 read parameters (28-10-10-90 cycles). The Cell Ranger package from 10X (25) was used to process the sequencing data using the *Drosophila* genome (dm6) and gene annotation downloaded from FlyBase (26) (release 6.29). Demultiplexing of DGRP strains to assigned timepoints was performed using demuxlet (12) as described in Dombrovski et al (15).

### Data Integration and clustering analysis

Between 200,000,000 and 500,000,000 reads per sample were processed with a minimum of 20 thousand reads per nuclei detected. The demultiplexing of DGRP genotypes also allowed us to identify cell doublets with a mean deduplication rate of 16-27%. After doublet removal, nuclei were filtered based on the following criteria: nuclei were removed with fewer than 175 or more than 3500 detected genes, fewer than 250 or more than 15000 total UMI and with percentage of mitochondria RNA greater than 10%. Following filtering, the mean unique transcripts (UMIs) across all the datasets was 3,246 per nuclei (median was 3,291) and the mean genes per nuclei was 951 (median was 1023).

We combined the nuclei with high confidence clock neurons from previous studies and performed integration and clustering analysis to identify clock neuron clusters (10, 18). Integration and clustering were performed in the Seurat package (v5) in RStudio (27). Nuclei and cells were split into separate layers and data was normalized, 2000 variable features were identified using FindVariableFeatures and these variable features were then used to integrate the datasets using IntegrateLayers and the Anchor-based CCA integration (27). The average cluster size for the LD dataset was 431 with a range from 191 (LPN_2 cluster) to 854 (LNd). For the DD dataset the average was 487 with a range from 162 (LPN_2) to 970 (LNd).

### Cycling transcript analysis

To identify cycling genes between circadian timepoints, we used the JTK algorithm of the MetaCycle package (28). We treated each DGRP line as a replicate and for the two timepoints where there were three replicates (ZT18 and ZT20), we averaged the values of two of the DGRP replicates. The average normalized TP10k value for each combination of cluster and ZT time point was calculated and used an input for cycling algorithms. We utilized two sets of cutoffs to be considered cycling, for loose the following cutoffs were used: a JTK_cycle *P* value of less than 0.05. For stringent determination, the following cutoffs were used: (i) a JTK_cycle *P* value of less than 0.05, (ii) a cycling amplitude (maximum expression divided by minimum expression) of at least twofold, and (iii) a maximal expression of at least 0.5 TP10K.

## Acknowledgements

We thank the members of the Rosbash lab for helpful discussions and critical reading of the manuscript. We greatly appreciate Dr. Yerbol Kurmangaliyev for insightful comments on the manuscript and for generously sharing fly stocks. This work was supported by the Howard Hughes Medical Institute (HHMI).

---

**S1:**
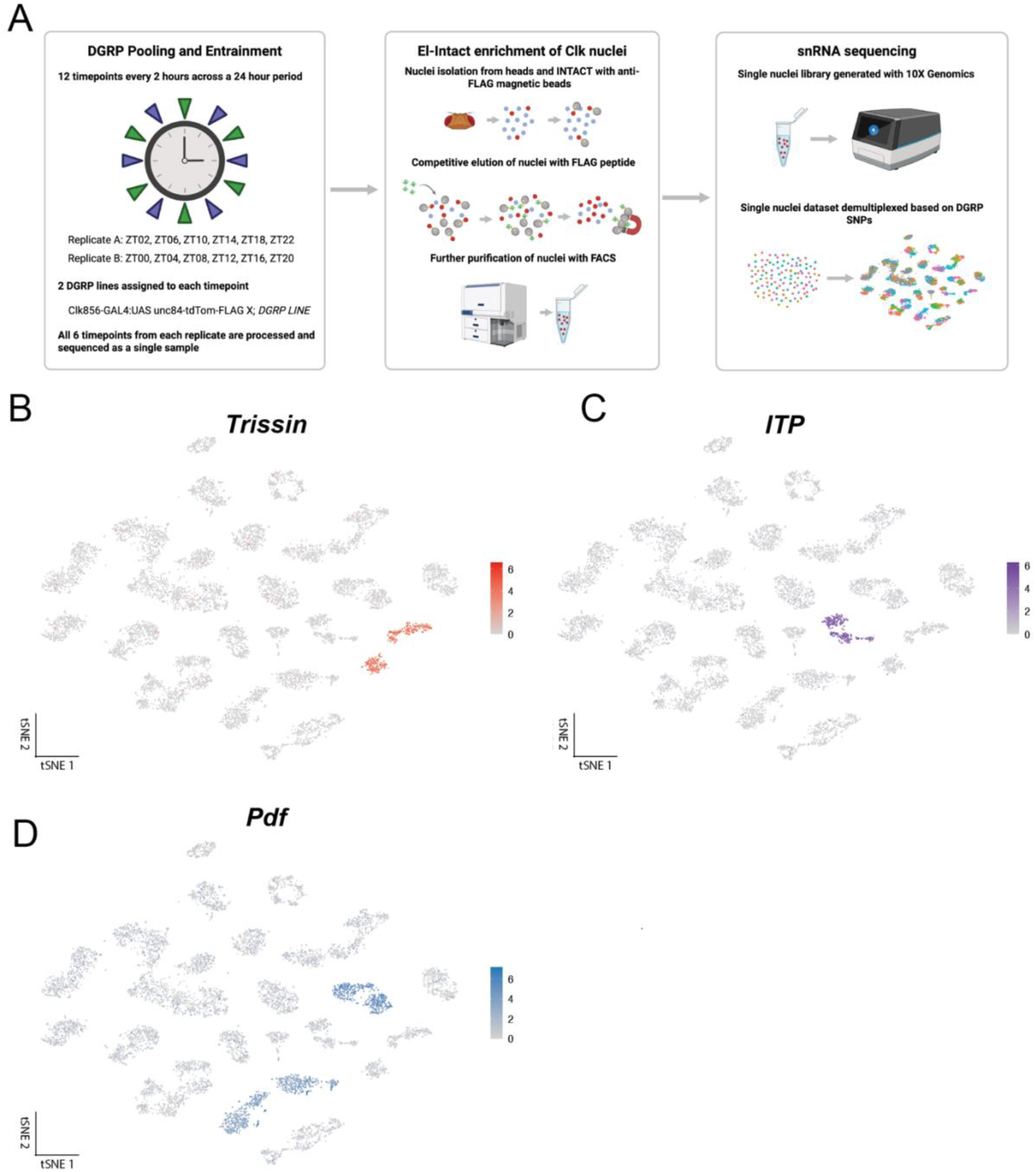
Methods of single-nuclei RNA sequencing of *Drosophila* clock neurons A) Schematic of experimental design for generating and processing single nuclei datasets using El-INTACT and pooled genetic multiplexing design. Flies with td-tomato-FLAG tagged labeled circadian neurons were collected across 12 time points, 2 hours apart split into two replicates of 6 time points. Td-tomato-tagged circadian nuclei were isolated using anti-FLAG magnetic beads, competitively eluted with FLAG peptide and purified with FACS. Single-nuclei RNA sequencing libraries were generated using single-cell 3’ RNA seq (10X Genomics). The schematic of the workflow was created using BioRender (https://biorender.com).T-SNE plots showing key lateral neuron marker genes expression: *Trissin* (B), *ITP* (C) and *Pdf* (D). (color bar, TP10K).

**S2:**
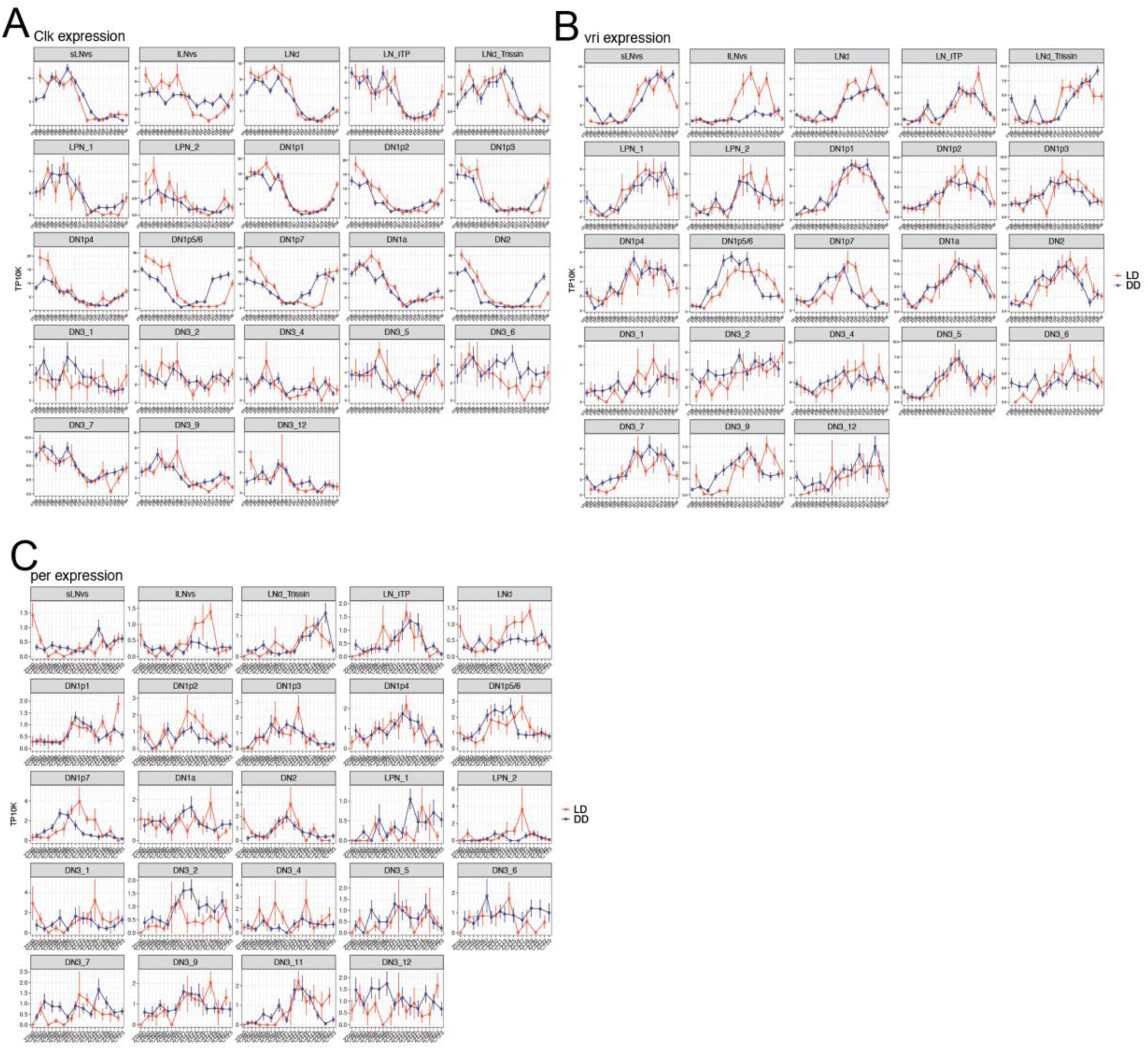
Expression of core clock genes in 12 time point snRNAseq (A-C) The mean *Clk* (A), *Vri* (B), and *per* (C) expression throughout the day in light: dark (LD) and constant darkness (DD) conditions is graphed for each cluster. X axis indicates the time points in LD and DD and Y axes are scaled independently for cluster. Error bars represent mean ± SEM. Red and blue lines indicate the gene expression in LD and DD conditions, respectively

**S3:**
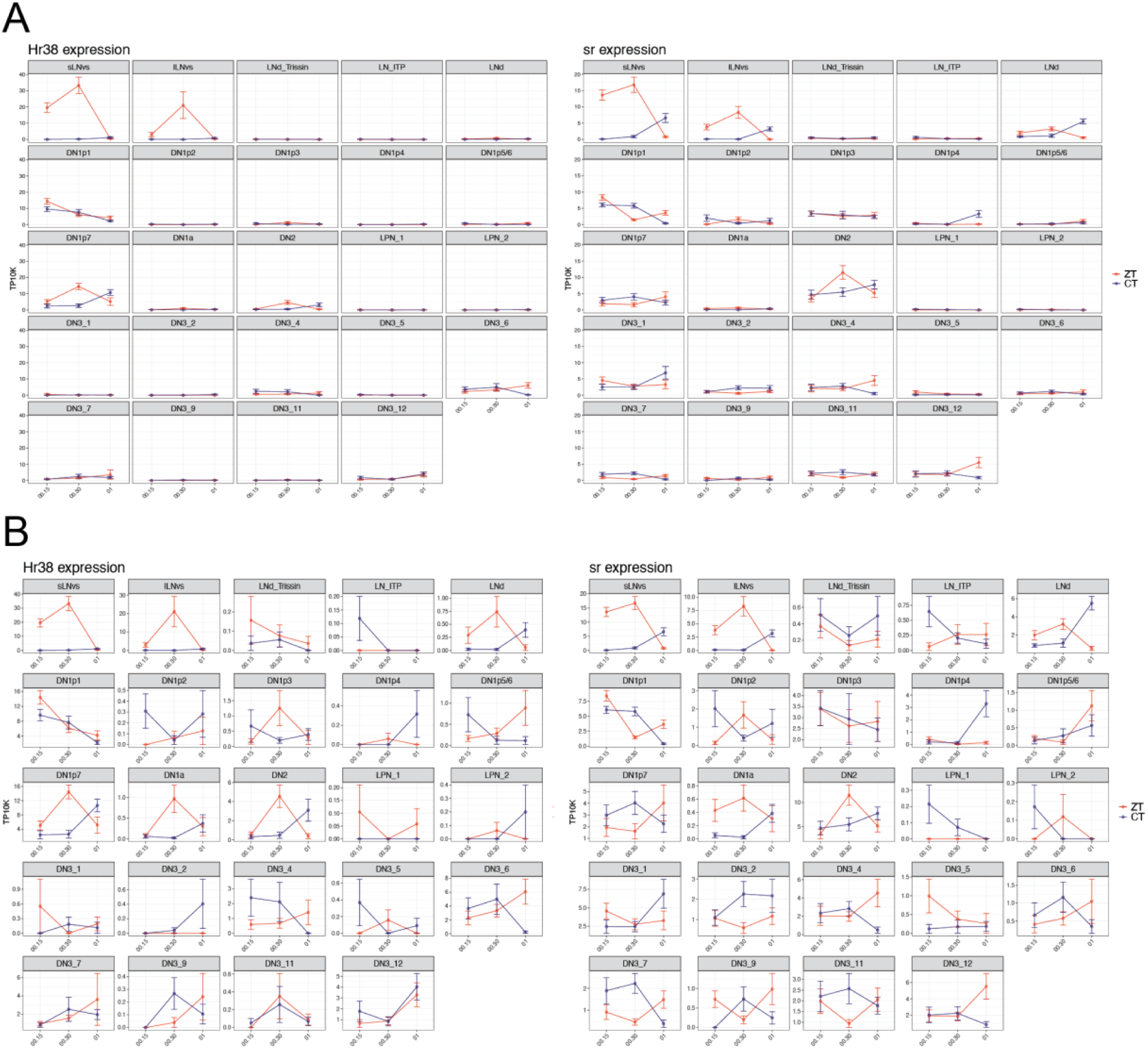
Hr38 and Sr show characteristic ARG expression with time course around lights on. A) The mean *Hr38* (left) and sr (right) expression following lights on at ZT00 in ZT conditions (red) or without light in the first day of DD (CT, blue). X axis indicates the time points following ZT00. Error bars represent mean ± SEM. B) *Hr38* and *Sr* expression of the same time course but with an independently scaled Y axis for each cluster.

**S4:**
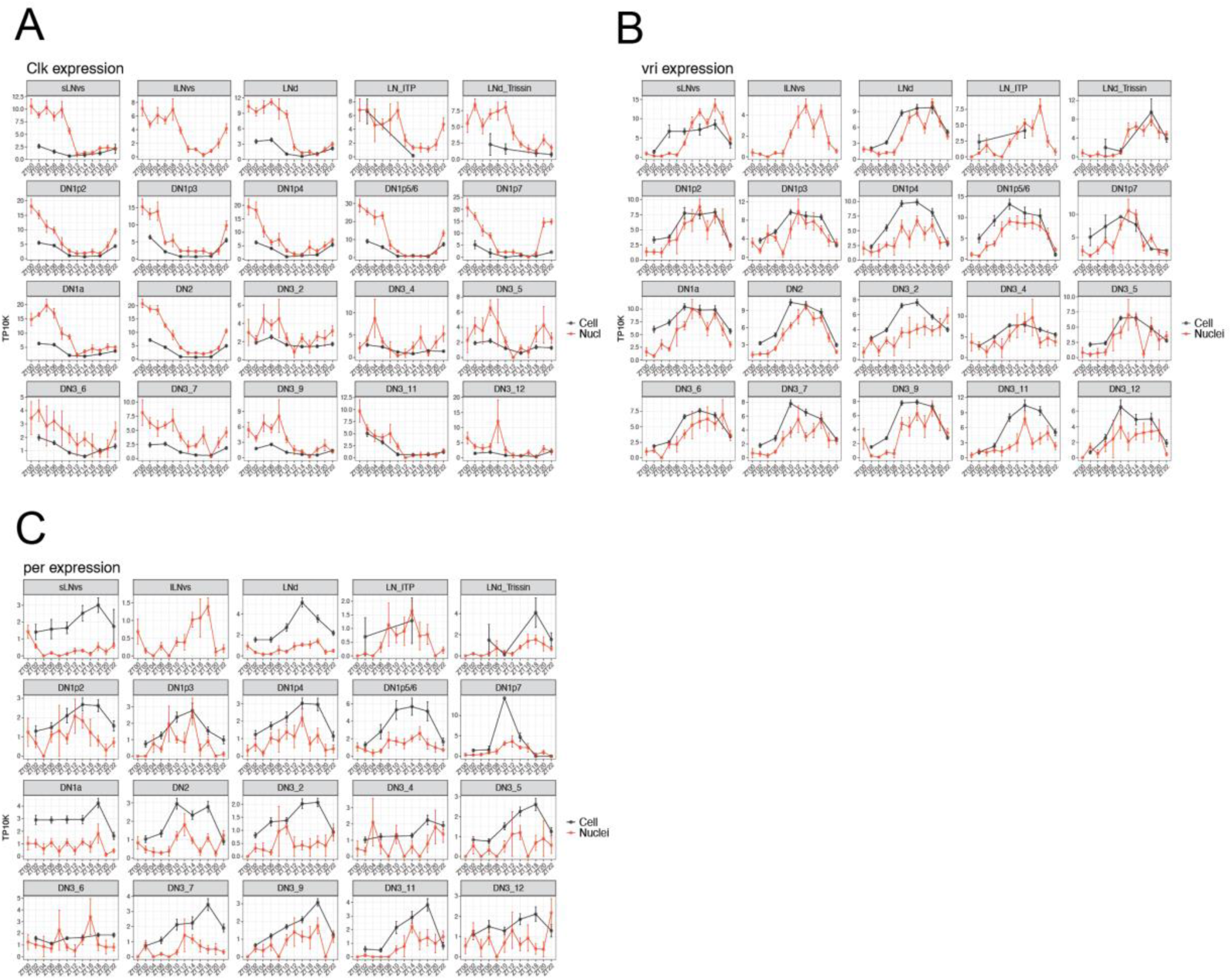
Expression of core clock genes in scRNAseq and snRNAseq (A-C) The mean *Clk* (A), *Vri* (B), and *per* (C) expression throughout the day in snRNAseq and scRNAseq datasets graphed for each cluster. X axis indicates the time points in LD and Y axes are scaled independently for cluster. Error bars represent mean ± SEM. Red and black lines indicate the gene expression in nuclei and cells, respectively.

**S5:**
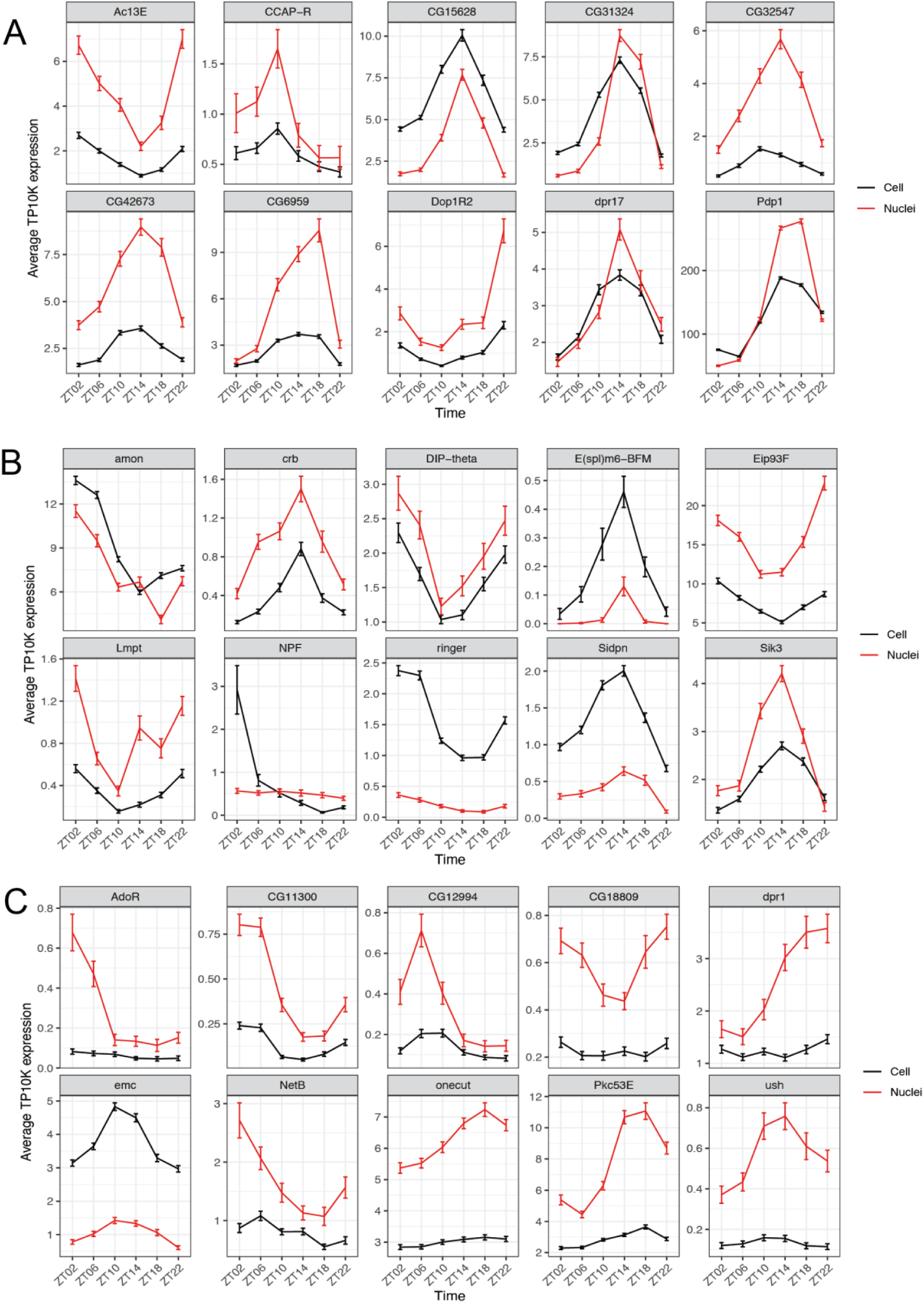
Pseduobulk expression of additional cycling transcripts in either cells, nuclei or both. A) Random selection of 10 genes from list of genes identified as cycling in both cell and nuclei dataset. B) Random selection of 10 genes from list of genes identified as uniquely cycling in cell dataset. C) Random selection of 10 genes from list of genes identified as uniquely cycling in nuclei dataset. Red and black lines indicate the gene expression in nuclei and cells, respectively. Y axes are scaled independently for each gene and error bars represent mean ± SEM.

